# Local olfactory nerve stimulation triggers patchy activation of columns across rat olfactory bulb

**DOI:** 10.1101/2025.01.29.635425

**Authors:** Israt Jahan, Lisa Kindler, Nicolas Reichardt, Hajime Suyama, Veronica Egger

## Abstract

Columnar organization is a hallmark of neocortical sensory areas, most prominently in primary somatosensory and visual cortex. In the olfactory bulb, the existence of a vertical columnar organization of circuits associated with glomerular input channels has been implicated by various approaches. Here, based on activity-dependent labelling and tissue clearing, we provide direct evidence for functional bulbar columns formed by mitral cells and granule cells that display a uniform morphology.

## Main Text

Similar to the neuroanatomical organization of mammalian neocortices and the retina, the olfactory bulb (OB) is a brain structure characterized by horizontal layering. Concurrently, the sensory input is segregated into glomerular input channels whereby each glomerulus receives convergent input from sensory neurons expressing the same olfactory receptor. The principal neurons connected to the same glomerulus, so-called ‘sister’ mitral cells (MCs)^1^, are by and large located below their respective glomerulus^2,3^, possibly indicating a vertical organization of the respective receptor channels. However, since the lateral dendrites of MCs extend horizontally over very long distances, a columnar arrangement of a given ensemble of sister MCs and their connected local interneurons such as parvalbumin neurons and granule cells (GCs) is not necessarily implied by the anatomy.

Evidence for a functional columnar organization of the OB involving GCs has first been derived from c-Fos expression data and voltage-sensitive dye imaging^4^, albeit at limited spatial resolution. Columnar structures linked to individual glomeruli were revealed by retrograde transsynaptic viral tracing^5,6^ and termed ‘glomerular unit columns’^7^; the mechanism underlying the apparently restricted spread of the virus might actually be activity-dependent^8^. Furthermore, there are functional indications for column-like clustering of GCs that respond to the same odorant during *in vivo* imaging, in particular for sparse activation^9^.

Here we present detailed evidence for functional columnar units based on activity-dependent labelling at high resolution upon local olfactory nerve stimulation in acute brain slices to mimic sensory input, using pERK as a marker of neuronal activity^10,11^. Stimulation was performed at medium depth of the olfactory nerve layer and is likely to activate segregated axonal fascicles of olfactory sensory neurons co-expressing the same receptor, so-called mesaxons^12,13^.

Local tetanic olfactory nerve stimulation within the medial olfactory nerve layer in horizontal acute brain slices of juvenile rats (see online methods) and subsequent staining for the activity marker pERK resulted in distributed columnar activation patterns of pERK-positive (pERK^+^) label ranging from the EPL to the deep GCL (Fig. 1A). The column closest to the stimulation site in the caudal direction also usually showed pERK^+^ signal in the glomerular layer. In order to facilitate the quantification of column dimensions, cell numbers and 3D reconstruction of columnar structures, we established brain clearing based on the CUBIC method^14^ in conjunction with pERK labelling (Fig. 1B, C; Supplementary movies S1, S2).

**Figure 1.**
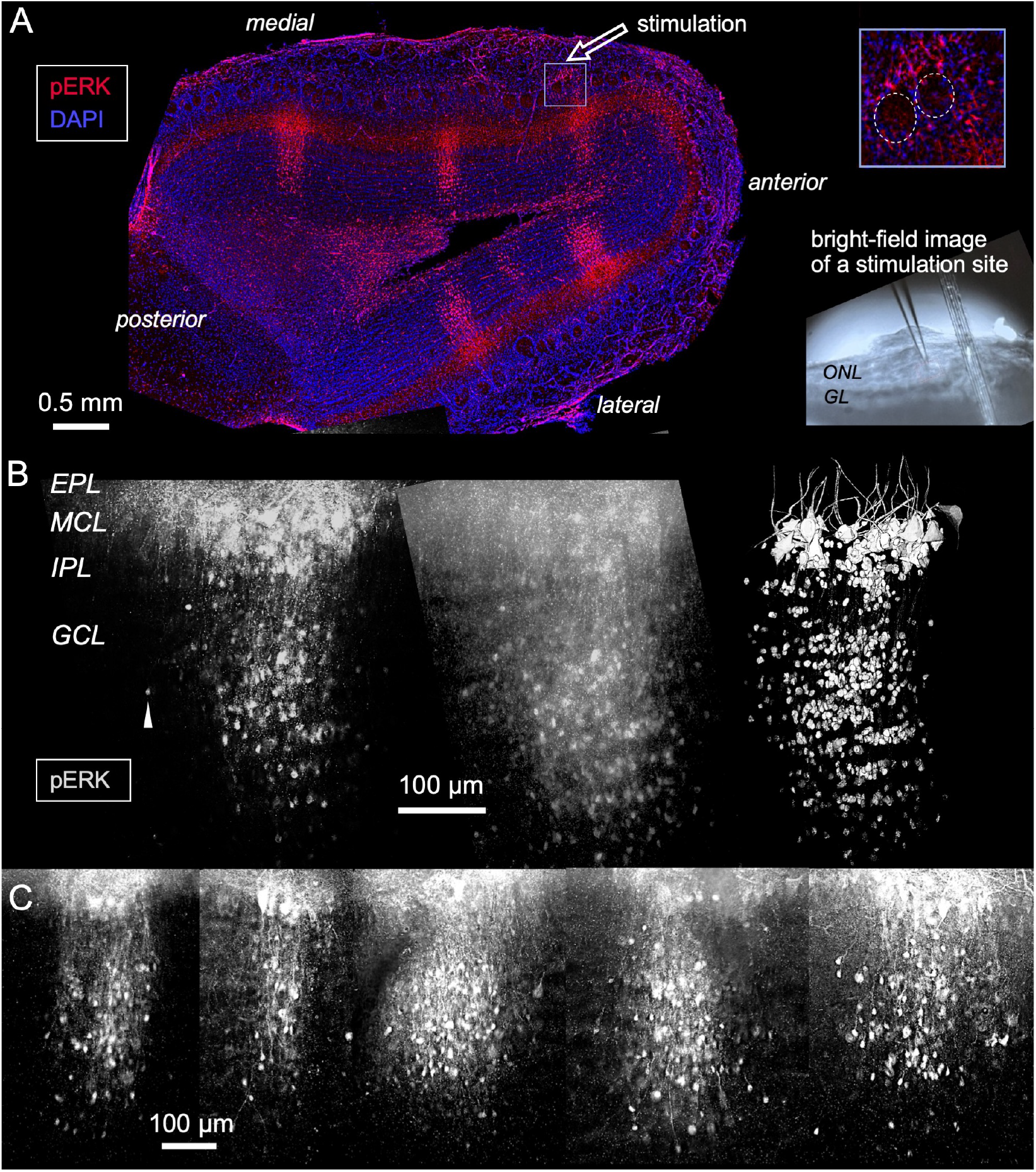
Columnar structures revealed by activity-dependent labelling. **A** Horizontal brain slice with stimulation site and multiple columns both on medial and lateral side (‘mirror columns’). Inset right: Magnification of area below stimulation site shows juxtaglomerular label (dashed line: glomerular insides). Bottom right: Wide-field image of stimulation site (in a different slice, grid right of the electrode). *ONL*, olfactory nerve layer, *GL*, glomerular layer **B** Representative column from cleared tissue. *EPL*, external plexiform layer, *MCL*, MC layer, *IPL*, internal plexiform layer, *GCL*, GC layer. Left: single 1 µm section of stack, arrow pointing to GC that was not counted as columnar. Middle: Maximum intensity z-projection. Right: 3D reconstruction (see Supplementary Movies S1, S2). **C** Selection of representative columns. All are images of 1 µm sections at largest horizontal extent of column. Same scale for all.

Tetanic ON stimulation was highly effective in triggering columnar activation since we observed columns in 42 of 52 stimulated slices (∼ 80 %). In 26 of the 42 slices with columnar activation, more than one column was labelled; frequently, additional columns were observed on the opposite (lateral) side of the slice, denoted as ‘mirror columns’ (in 17 out of the 26 slices with at least 2 columns, see Fig. 1A); such mirror columns were never observed in isolation, i.e., without any columns on the medial side. Independent of the location of a column relative to the stimulation site, pERK^+^ cell bodies were observed mostly in the MCL and GCL, with only few cells in the EPL and IPL. Unstimulated control slices stained for pERK (n = 19) did never show columns.

To rule out overstimulation by the applied tetanus as a cause of columnar activation, we also performed a set of experiments using theta burst stimulation (TBS). Theta bursts are central to olfactory processing since the sensory activation of MCs occurs in gamma bursts that are spaced by the respiratory theta rhythm^15,16^. TBS was similarly effective in columnar labelling (in 67% of 24 stimulated slices), and the resulting columns did not differ significantly in all the analysed parameters except for a tendency for less pERK^+^ MCs upon TBS (Mann-Whitney tests, Supplementary Table).

Therefore data from cleared tissue were pooled for the cumulative analyses of columnar morphology shown in Figure 2A (n = 31 columns): The mean height of columns y was 488 ± 78 µm (all means ± S.D.), and their diameter in the slice plane, the width x, was 267 ± 72 µm, similar to the mean depth of columns in the slice z, 253 ± 77 µm (shrinkage-corrected, see Methods). While most columns had oval cross-sections as evident from the z/x ratio, the average cross-section was circular (mean log_2_(z/x) = −0.09 ± 0.53), indicating that there is no preferred orientation of columnar cross-section shape with respect to the horizontal OB slice plane. The mean cell count of columnar MCs was 11 ± 5, in line with earlier observations on ‘sister MCs’ belonging to the same glomerulus^17,18,19,20^. The mean columnar GC count was 146 ± 57. This number lies below the number of GCs per glomerulus in juvenile rat (total OB GC number 2 million^21^ divided by number of glomeruli 4000^17^= 500 GCs). Possibly, since tufted cells seem not to participate in the columnar architecture revealed here, their GC partners are also not involved. Moreover, this result might imply that any GC is unlikely to belong to more than one column, at least in terms of its activation sufficient for pERK expression, even though a GC can obviously connect to MCs belonging to many different columns. All these parameter distributions were unimodal and close to normal; thus, columns appear fairly stereotyped. Also, the density of GCs within the estimated columnar volume (cross-section • height z) was mostly highly homogenous (Fig. 2A, mean 6.5•10^-6^ ± 3.5•10^-6^ µm^-3^, median 5.4•10^-6^ µm^-3^).

**Figure 2.**
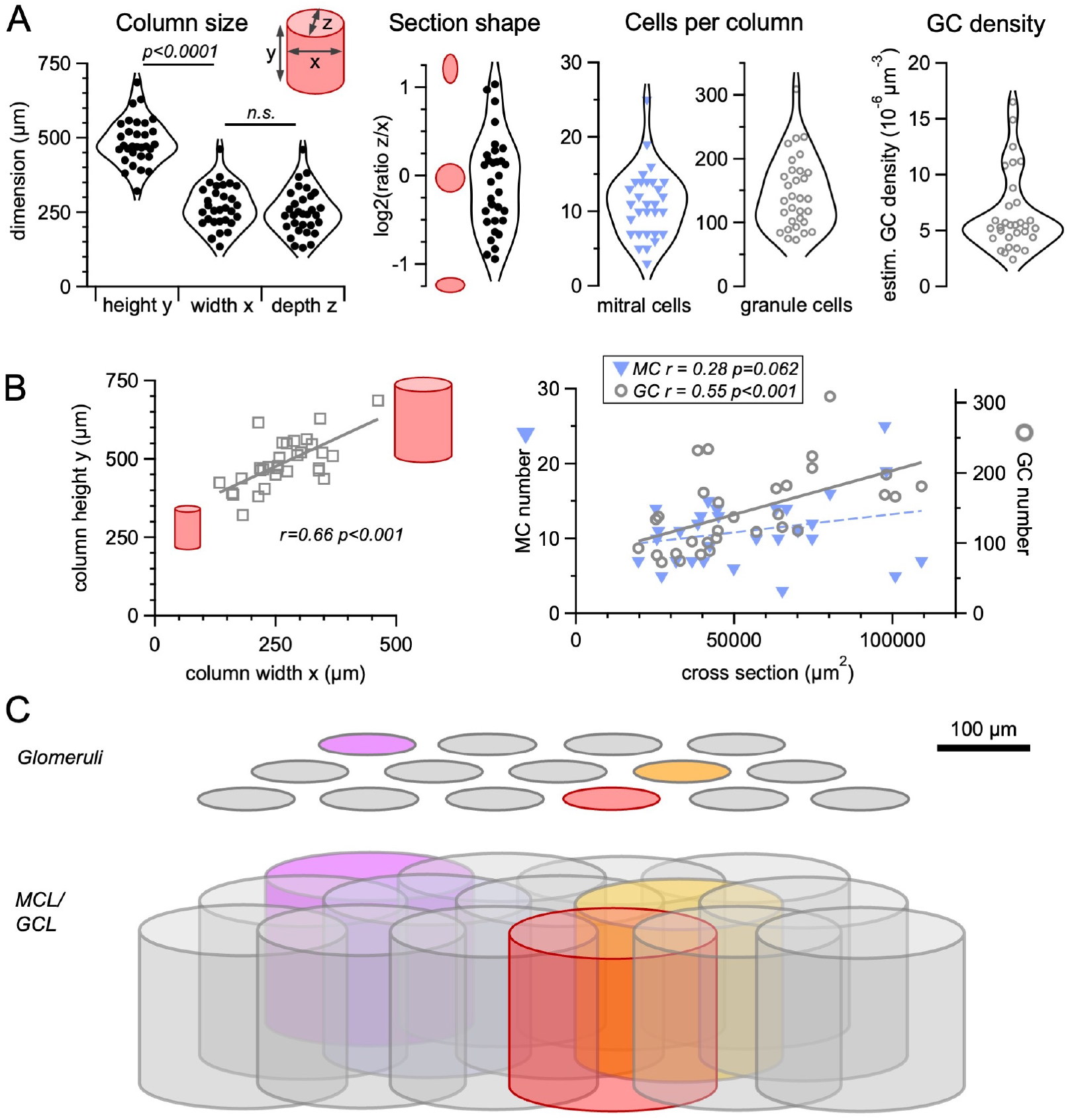
Cumulative analysis of columns (n = 31) See Supplementary Table for statistics. **A** Distributions: Columnar dimensions y, x, z; Elliptic shape of the columnar cross-section described by log_2_(z/x). 0: circular cross section z = x, 1: ellipse with z = 2x, −1: ellipse with z = x/2; Cell counts per column (MCs, GCs); GC density (see Methods). Mean values ± standard deviations given in main text. **B** Correlations: Left: height y vs width x. Right: MC count and GC count versus columnar cross-section. **C** Schematic depiction of lateral extent of glomeruli and lateral extent of their associated columns. Y-dimension not to scale, elliptic shape of columns not accounted for. Colored glomeruli and columns symbolize activation by an odor.

Figure 2B shows that the height of columns was significantly correlated with their width, with taller columns having a larger diameter, and the cross-section of columns positively correlated with the GC count, whereas for the MC count there was a weak tendency. These relations are well in line with the fairly uniform estimated density of columnar GCs.

The subset of analysed mirror columns (n = 7 out of the 31 analysed columns) did not differ significantly from non-mirror columns in any of these parameters (Supplementary Table). Therefore, we conclude that the observed columns must be organized by a similar type of excitatory connectivity, independent of their location with respect to the mid-axis. While multiple medially located columns might be due to stimulation of several segregated fascicles of the ON, the nature of this connectivity for the mirror columns is unknown at this point. Possibly it is related to the columnar intrabulbar association system between the medial and lateral glomeruli innervated by the same type of olfactory receptor neuron, in which a subtype of superficial tufted cell projects to GCs of the homologous column (e.g. ^22,23^). However, this system is most likely not yet fully established in juvenile rats^24^ and also cannot explain the labelling of MCs in mirror columns at this point.

A functional columnar clustering of GCs below activated MCs was predicted by computational models^25,26^ and could facilitate synchronization of the MCs belonging to coactive glomerular channels^25^. Such clusters might form through activity-dependent learning^27^. While the columns described here might not be identical to the columnar structures revealed by viral tracing^5^, we would like to suggest that there is at least a partial homology. In any case, our confirmation of activity-dependent MC-GC columns might inform updates of detailed network models of the bulb (e.g. ^28,29,30^) and help to further dissociate the function of GCs from that of other bulbar inhibitory interneurons such as periglomerular cell types and Parvalbumin neurons.

Another important consequence of our study is that columns must be substantially overlapping in space as illustrated in Figure 2C, since their mean diameter of almost 300 µm exceeds the mean diameter of glomeruli (∼120 µm in rat,^17^). Thus bulbar columns are ‘less rigidly defined’^4^ than cortical columns. This spread is in line with and possibly implemented by the lateral spread of sister MC somata on the order of 200 µm^2,3,20^. It also fits with our model of isotropic connectivity strictly based on known anatomical parameters^31^, whereby MC-GC connectivity is estimated to peak within a radius of 100 µm around a given MC and the probability of a GC to connect to two sister MCs is also highest within a similar regime. It remains to be elucidated whether there is also functional overlap for nearby columns activated by the same odor.

Functional columnar structures thus emerge as an overarching principle of neural organization beyond the neocortex, both in non-cortical areas in mammals such as the OB and also in non-mammalian species^32^.

## Author contributions

IJ designed research, performed experiments, analyzed data, drafted Methods section.

LK performed experiments.

NR performed experiments.

HS supervised research.

VE conceived and designed research, analyzed data, prepared figures, drafted manuscript.

All authors approved the final version of the manuscript.

## Material and methods

### Animals

Both WT and VP-eGFP^33^ juvenile rats aged PND12-19 of both sexes were used, in line with our 3R policy; the use of different strains should also increase the significance of results (see^34^). For the brain clearing and quantitative analyses, Wistar rats (n = 4) and VP-eGFP rats (n = 6) aged PND15-18 were used. The data sets derived from WT and VP-eGFP rats did not show any statistical differences (n = 14 and n = 17 columns, respectively, see Supplementary Table). The rats were bred in the animal facility of the University of Regensburg and housed under standardized laboratory conditions (temperature: 20-26^°^C, humidity: 60-65%, 12/12 light dark cycle and food ad libitum).

### Preparation of olfactory bulb slices

First, artificial cerebrospinal fluid (ACSF: 125mM NaCl, 26mM NaHCO3, 1,25mM NaH2PO4, 20mM Glucose, 2,5mM KCl, 1mM MgCl2, und 2mM CaCl2) was incubated on ice and bubbled with carbogen (95% O_2_ and 5% CO_2_) for 30 minutes to stabilize pH and oxygen levels. Rats were deeply anesthetized with 100% isoflurane and decapitated. All experiments were performed in compliance with national and institutional regulations for the care and use of laboratory animals, following the guidelines set by the EC Council Directive (86/89/ECC) and German animal welfare laws^10^. Horizontal slices (thickness 300 or 500 µm) were prepared using a Vibratome (Leica 1200 VT), with the tissue being immersed in ice-cold ACSF and with continuous supply of carbogen. OB slices were then incubated in ACSF at 37°C for 30 minutes. The slices were incubated for further 3 hours at room temperature before the stimulation experiment to prevent pERK signals due to initial spontaneous activity^35^. For the controls without any olfactory nerve stimulation, we used OB slices after either 30 minutes of incubation at 37°C (n = 6) or after further 3 hours of incubation at room temperature (n = 13).

### Olfactory bulb stimulation

For stimulation, slices were placed in a chamber under a light microscope (Axioplan, Zeiss, Germany) and perfused with carbogenized ACSF (see above). A glass capillary micropipette with a fine tip (sized approx. 2 MOhm) was pulled (Narishige, Tokyo, Japan). The micropipette was filled with ACSF and a stimulation electrode was inserted into it. The stimulation electrode was connected to an external stimulator (STG 1004, Multichannel systems, Reutlingen, Germany) controlled by a PC. A stimulation location was then selected in the anterior middle part of the olfactory nerve layer on the medial part of the OB (see Fig. 1A), and the stimulation electrode was carefully lowered onto the pre-determined location until it visibly penetrated the tissue. Finally, the slice was stimulated, using Multichannel systems software (V2.1.5). The strength of stimulation was 800 µA for 100 µs. Tetanic stimulation was performed at 50 Hz for 1 min, i.e. 300 stimulations in total; theta burst stimulation (TBS)^36^ involved a sequence of five bursts of five stimulations at 50 Hz, spaced 5 Hz apart, and repeated ten times at an interval of 10 s (250 stimulations in total). A picture was also taken at this point to record the stimulation location. After 5 minutes the stimulation electrode was slowly removed, and the slice was kept in ACSF for 5 more minutes. Next, the slice was transferred to a 12 well plate and incubated in 4% paraformaldehyde (PFA-PBS) overnight at room temperature for fixation.

### Initial set of experiments

In an initial set of experiments without tissue clearing (not used for the quantitative analyses), 300 µm OB slices were resectioned into 30 µm slices and underwent immunohistochemistry against pERK, including DAPI staining (for details see^10,11^).

### Tissue clearing

We followed the CUBIC protocol^14^ for tissue clearing, using two reagents, ScaleCUBIC-1 (Reagent-1) and ScaleCUBIC-2 (Reagent-2, see Table for ingredients).

**Table 1:**
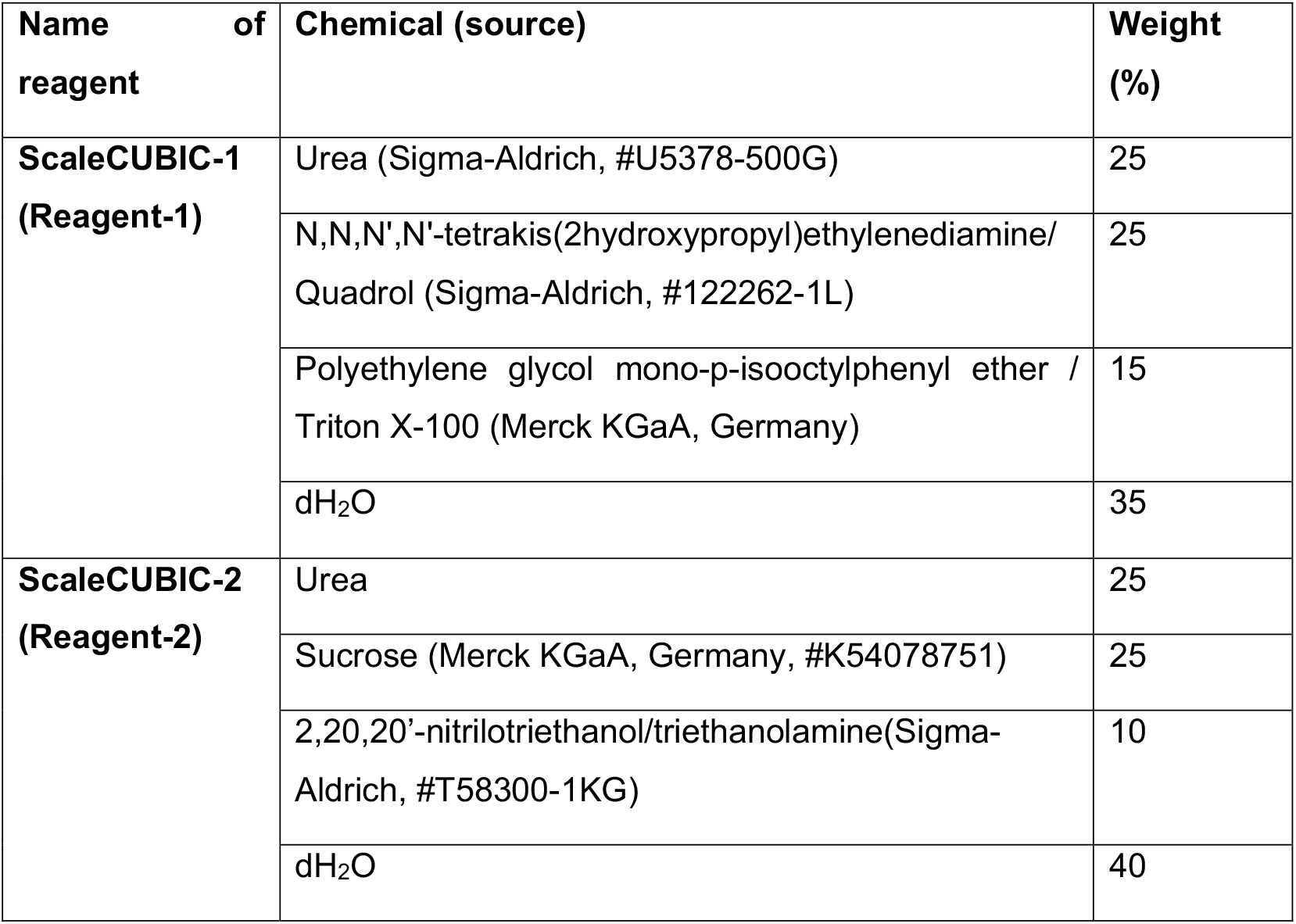
List of chemicals used for CUBIC tissue clearing protocol (from^14^)

| Name of reagent | Chemical (source) | Weight (%) |
| --- | --- | --- |
| **ScaleCUBIC-1 (Reagent-1)** | Urea (Sigma-Aldrich, #U5378-500G) | 25 |
|  | N,N,N',N'-tetrakis(2hydroxypropyl)ethylenediamine/Quadrol (Sigma-Aldrich, #122262-1L) | 25 |
|  | Polyethylene glycol mono-p-isooctylphenyl ether / Triton X-100 (Merck KGaA, Germany) | 15 |
|  | dH<sub>2</sub>O | 35 |
| **ScaleCUBIC-2 (Reagent-2)** | Urea | 25 |
|  | Sucrose (Merck KGaA, Germany, #K54078751) | 25 |
|  | 2,20,20'-nitrilotriethanol/triethanolamine(Sigma-Aldrich, #T58300-1KG) | 10 |
|  | dH<sub>2</sub>O | 40 |

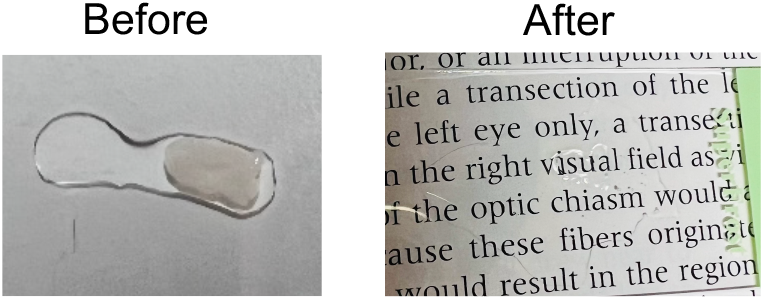

Following overnight fixation with PFA, the OB slices were either incubated in PBS at 4°C for 2 days with daily solution changes or washed two times with PBS at room temperature on a shaker for 2 hours each. Subsequently, the slices were transferred to falcon tubes and incubated in diluted Reagent-1 (1:1 in dH_2_O, 2ml per slice) at 37°C for 4-6 hours inside a thermal mixer (Eppendorf ThermoMixer Comfort 6901, Hamburg, Germany), featuring a heating block designed for 4 x 50 ml Falcon tubes (shaking speed 300 rpm). Afterward, the diluted Reagent-1 was replaced with Reagent-1 (∼5 ml per tube), and the slices were incubated overnight at 37°C. This process was repeated for another four days for 500 µm slices. To clear 300 µm slices, 3 days of incubation with Reagent-1 was sufficient. The Reagent-1 solution was changed every two days. Immunohistochemical staining (see next section) was performed after completion of incubation with Reagent-1. After that, the slices were immersed in Reagent-2 overnight at 37°C. The following day, the Reagent-2 solution was exchanged, and the slices were again incubated at 37°C in Reagent-2 for approximately 24 hours.

### Immunohistochemistry protocol

All immunohistochemistry was carried out at 37°C and 300 rpm in the thermal mixer. Immunohistochemical solutions were prepared according to^10^. After completing incubation with Reagent-1, OB slices were washed with PBS 3 times for 2 hours each. Approximately 2 ml of PBS was used for each washing step. Then, the tissue was incubated in 0.1M glycine in PBS for 20 minutes, followed by a 10-minute wash with PBST. Next, the tissue was incubated in a blocking solution of 5% normal donkey serum (NDS, NB-23-00183-1, NeoBiotech, Nanterre, France) in 0.2% cold water fish skin gelatin (AURION, Wegeningen, Netherlands) and 0.01% NaN_3_ for 1 hour. Next, the tissue was further incubated with the primary antibody (1 ml) for 72h. Subsequently, the tissue was washed with PBST 4-6 times for 10 minutes each. Finally, the tissue was incubated with the secondary antibody (1 ml) for 48h. The OB slices underwent another 4-6 washing steps with PBST for 10 minutes each before applying the Reagent-2 solution (see above).

### Mounting and imaging

After completing the incubation with Reagent-2, the slices were gently transferred to an objective slide (Thermo Fisher Scientific, Waltham, MA USA) and mounted with Mowiol (matching refractive index, 1ml, Sigma-Aldrich, #STBG2469V). Following mounting, the slides were left in the dark for drying at room temperature overnight. Next, the slices were imaged using a DM6 B microscope from LEICA, equipped with LAS X software. An excitation wavelength of 594 nm (for pERK) was employed. The exposure time was set to 500 ms. A z-stack was captured, with each z-section spaced 1 μm apart. The acquired image was further processed using the “Lightning & Thunder Processing” tool.

### Image analysis

To analyze the images (in .lif format), Fiji/ImageJ was employed. The Fiji/ImageJ software contains a built-in tool named “Cell Counter”, which allows for the selection of cells throughout a z-stack. The check box for “show all” should be selected to prevent multiple selection of the same cell. Before counting, column borders were delineated by eye, separating single isolated pERK^+^ GCs from clouds of pERK^+^ GCs with intercell distances ≲40 µm^5^.

Column dimensions (Fig. 2A right inset) were estimated as follows: For the height z of the column, a vertical line was drawn from the MCL to the bottom end of the GC column, and the length of this line was measured using the measure function in Fiji/ImageJ. The width x of the column was measured in the same way by drawing a parallel line to the MCL at the level of the broadest part of the column in the GCL. For the depth z (the diameter along the z-axis of the stack) the number of z-sections (1 µm each) between the first appearance of pERK^+^ columnar GCs and their disappearance was multiplied by a factor of 2 to correct for shrinkage^37^. The density of GCs in a given column was estimated by dividing the GC count by the columnar volume which in turn was estimated as the cross-sectional area π•(x/2)•(z/2) times the height y. Note that this measure is likely to overestimate the columnar volume, since x, y, z were determined as the maximal extent of the column in either direction.

### 3D reconstruction

First, the z-stack was imported into Fiji/ImageJ. To ensure a consistent analysis of all sections, the brightness and contrast were adjusted to set the maximum value to 2700 using the “Brightness & Contrast” function, resulting in a standardized maximum range of 30 to 2700 for all analyzed files. The z-stack was then separated using the “stack to images” function, and the images were sequentially saved as .tiff files. The concatenated images containing focused pERK^+^ GC somata were retained for further analysis, while the rest were discarded.

Subsequently, this reduced z-stack was loaded into Fiji/ImageJ’s TrekEM2 (Blank) tool. The image properties (pixel width, pixel height, voxel depth) were also corrected to match the original image. To rectify z-shrinkage due to the mounting process, the z-extent of the stack was normalized according to the original thickness of the slice. After that, GC and MC somata as well as GC and MC apical dendrites were selected in each image using the “selection tool”. Once all cells in each image had been selected, the selected area was extracted using the “Export-Image stack under selected area” option. Finally, the “3D Viewer” plugin was used to generate a 3D image of the reconstructed activated column. The movie of its 360-degree rotation was generated using the Path View > Record 360° Rotation feature. The file was then saved in AVI format at a rate of 12 frames per second.

### Data analysis

The cumulative data were plotted in IGOR64 (Wavemetrics, Oregon, US). All statistics are based on the non-parametric Mann-Whitney or Wilcoxon tests (with the outcomes listed in the Supplementary Table) and the basic correlation statistics available on http://vassarstats.net/. Mean values are given with standard deviation (S.D.).

## Supplementary Material

Supplementary Table below

**Supplementary Table.**
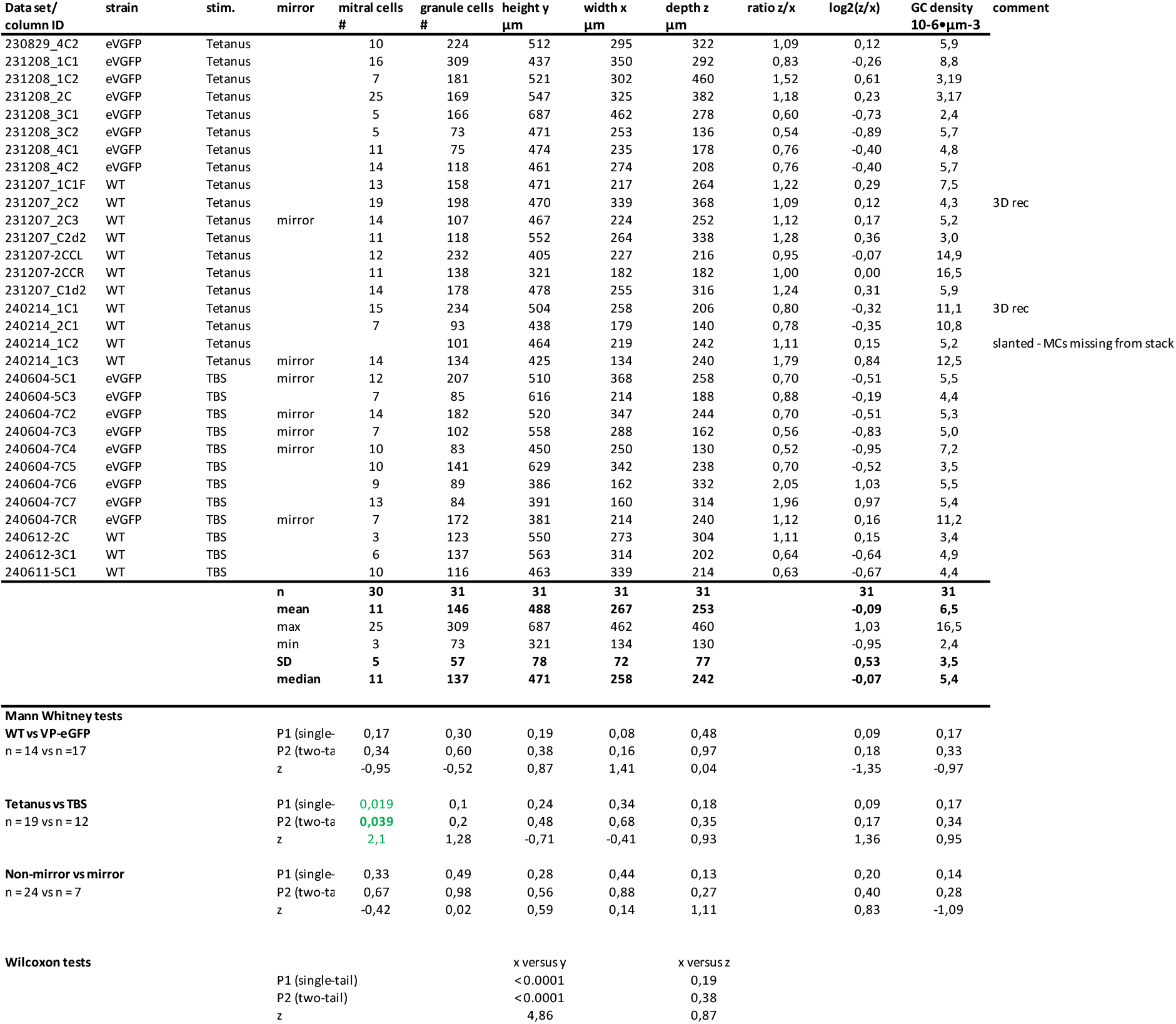

Supplementary Movies S1, S2 at

Mendeley Data

https://data.mendeley.com/preview/t93hjt6wg3?a=d97a3739-5222-4366-b45a-5fb42a212e6e

Reserved DOI:10.17632/t93hjt6wg3.1

